# Shared and distinct default mode systems in mental time travel and affective experience over time

**DOI:** 10.64898/2026.01.20.700521

**Authors:** Yuan Zhang, Wenyi Dong, Kun Fu, Menghan Zhou, Yanan Qin, Raymond C.K. Chan, Keith M Kendrick, Dezhong Yao, Shuxia Yao, Benjamin Becker

## Abstract

Overarching conceptualizations propose a critical role of the default mode network (DMN) in self-referential mental time travel, particularly in autobiographical memory retrieval and episodic future thinking, and internal (intrinsic) emotion generation and regulation. However, these conceptualizations have not been directly evaluated. Against this background, the present fMRI study aimed to identify both shared and distinct neural systems underlying autobiographical episodic processing across different temporal contexts - specifically, episodic memory retrieval (EMR) and episodic future thinking (EFT) - and to examine how these systems interact with affective experiences, including valence and arousal. Our findings demonstrated the central role of the DMN - encompassing the medial prefrontal cortex (mPFC), posterior cingulate cortex (PCC), and medial temporal lobe (MTL) - in both EMR and EFT. Importantly, we identified a functional dissociation along both valence and temporal dimensions: the ventromedial prefrontal cortex (vmPFC) was more strongly associated with positive experiences and simulations, whereas the dorsomedial prefrontal cortex (dmPFC) was consistently engaged during the processing of negative affect across past and future contexts. Moreover, representational similarity and parametric analyses indicated that the hippocampus supports differential processing of valence and arousal across temporal domains. Together, these findings provide empirical evidence for the involvement of cortical midline core DMN systems in autobiographical processing across time and suggest overlapping and distinct systems for the integration of emotional experiences across mental time travel.

## Introduction

Mental time travel, the human capacity to mentally reconstruct affective experiences from the past (episodic memory retrieval, EMR) and to simulate potential affective events in the future (episodic future thinking, EFT). Accumulating evidence suggests that this dual temporal projection system enables not only autobiographical continuity but also goal-directed planning, affective forecasting, and counterfactual reasoning—cognitive functions critical for survival and social functioning [1–3]. Convergent evidence from neuropsychology and behavioral economics highlights its pivotal role in decision-making processes, where impaired mental time travel correlates with myopic choices and reduced hedonic optimization in healthy populations [4,5]. In clinical populations, the disability in mental travel is closely associated with mood disorders, such as excessive negative prospecting in depression and generalized anxiety [6,7], maladaptive EMR patterns underlie intrusive negative memory recall in post-traumatic stress disorder (PTSD) [8–10] and overgeneralized autobiographical memory in borderline personality disorder [11,12].

The “constructive episodic simulation hypothesis” suggests that both processes rely on a common neural system for recombining episodic details, yet EFT demands additional mechanisms for novelty generation and uncertainty resolution [13]. Convergent neuroimaging evidence suggests that these processes rely on a shared neural substrate—the default mode network (DMN)—encompassing the hippocampus, medial prefrontal cortex (mPFC), posterior cingulate cortex (PCC), and angular gyrus [2,14,15]. However, emerging evidence also showed inconsistent differences between these two processes. For instance, the EFT induced stronger engagement in the lateral and medial parietal cortex and hippocampus [15–17], whereas EMR induced greater activation in the visual regions [18] and even the hippocampus [19,20]. These differential activation patterns suggest that temporal directionality modulates neural resource allocation, though the mechanisms remain rarely explored.

While accumulating evidence suggests a role of the DMN in emotion generation and regulation [21] and in everyday life a considerable amount of emotional experience is elicited by internal processes, including mental simulation and autobiographical mental time travel [22,23], research on the neural basis of emotions has strongly focused on emotional experiences induced by external stimuli, including affective pictures or movies (e.g., [24–26]). While initial studies have examined the interaction between autobiographical mental time travel in terms of construction and elaboration of emotional experiences [18], of focusing on temporal context within the same valences [23,27], or social versus non-social context within a single temporal domain [28], the interaction between basic emotional domains such a valence and arousal within the EMR-EFT framework has not been systematically determined. Initial findings suggest that emotion modulates hippocampal-prefrontal coupling during positive autobiographical recall [29], while negative future simulations amplify amygdala-insula connectivity [18]. Moreover, recent articles indicated the core network, the DMN, also serves a critical role in affective and cognitive processes [30–32], such as self-evaluative processing, social perspective-taking, episodic memory and foresight. Yet there is not enough evidence to systematically examine how valence interacts with temporal direction (past vs. future) to alter large-scale network dynamics, which is a critical gap given the clinical relevance of valence-biased mental time travel in mood disorders.

Against this background, the aim of the present study is to identify shared and distinct neural systems underlying the EMR and EFT through a 2×3 factorial fMRI design (details see Fig 1) manipulating temporal context (EMR vs. EFT) and emotional valence (positive, neutral, vs. negative). Based on previous findings, we hypothesized that (a) the EMR and EFT tasks would exhibit shared DMN activation across all valence conditions, particularly in the mPFC and PCC nodes supporting self-projection, and (b) distinct neural signatures would emerge in visual cortices (EMR>EFT) and frontoparietal regions (EFT> EMR) through univariate general linear model (GLM) analysis and it may reveal some key regions encoding differences of interaction of temporal and valence conditions through multivariate pattern analysis (i.e. representational similarity analysis; RSA). Thus, this study aimed to advance theoretical models of episodic cognition while providing neural targets for interventions addressing maladaptive mental time travel in psychopathology by elucidating how temporal and affective dimensions jointly shape mental time travel.

**Fig 1.**
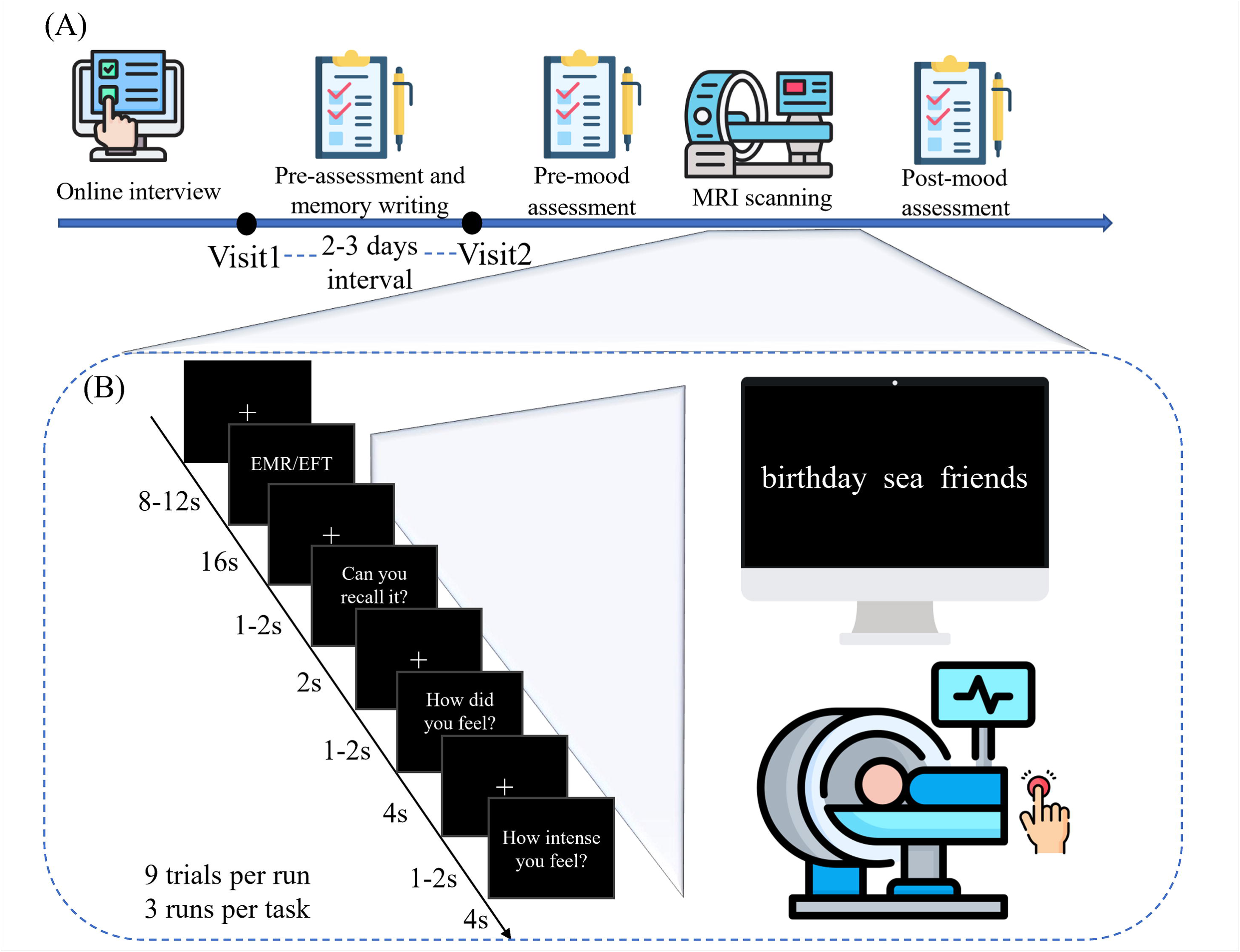
Experimental protocol. **A** Experimental procedure. Participants first completed questionnaires and vividly imagined and documented positive, neutral, and negative affective events from the past (episodic memory recall; EMR) or imagined in the future (episodic future thinking; EFT). During the subsequent fMRI session during the second visit, participants underwent an anatomical T1-weighted scan, resting-state fMRI acquisitions, and EMR and EFT tasks. **B** Task paradigm. Each trial began with an 8-12s jittered fixation cross, and 3 cue words (based on the personalized documentation from the first day) were displayed on the screen to instruct participants to perform the EMR or EFT task. During this period particiants were asked to use button presses to indicate the duration of the memory or imagination, respectively. Following each trial participants reported their subjective experience of valence and arousal for this event.

## Results

### 2.1 Behavioral results

Paired-sample t-tests on mood changes were evaluated by examining the PANAS scores before and after MRI. The results showed no significant differences in the negative subscale (t=1.508, *p*=0.138, Cohen’s d=0.21), but significantly decreased scores in the positive subscale after EMR and EFT tasks compared to before tasks (t=2.942, *p*=0.005, Cohen’s d=0.41).

We conducted a repeated-measures ANOVA on valence, arousal, and vividness scores with valence and temporal context conditions as within-subject factors. For valence scores, a significant main effect of valence (F(2,102)=590.573, *p*<0.001, 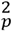=0.921) and a significant main effect of temporal context (F(1,51)=12.577, *p*=0.001, 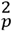=0.198) was observed in the context of a significant valence×temporal context interaction (F(2,102)=10.089, *p*<0.001, 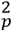=0.165; Fig 2A). Post-hoc analyses demonstrated significantly higher valence ratings in the positive condition relative to both neutral and negative conditions across past and future scenarios (all *ps*<0.001; details see S1 Table), and lower ratings for the negative events relative to the neutral ones for past and future scenarios (all *ps*<0.001). Significantly lower scores for valence ratings in the past relative to the future were only found in the negative condition (*p*<0.001), indicating a more negative feeling for negative events in the past than the future.

**Fig 2.**
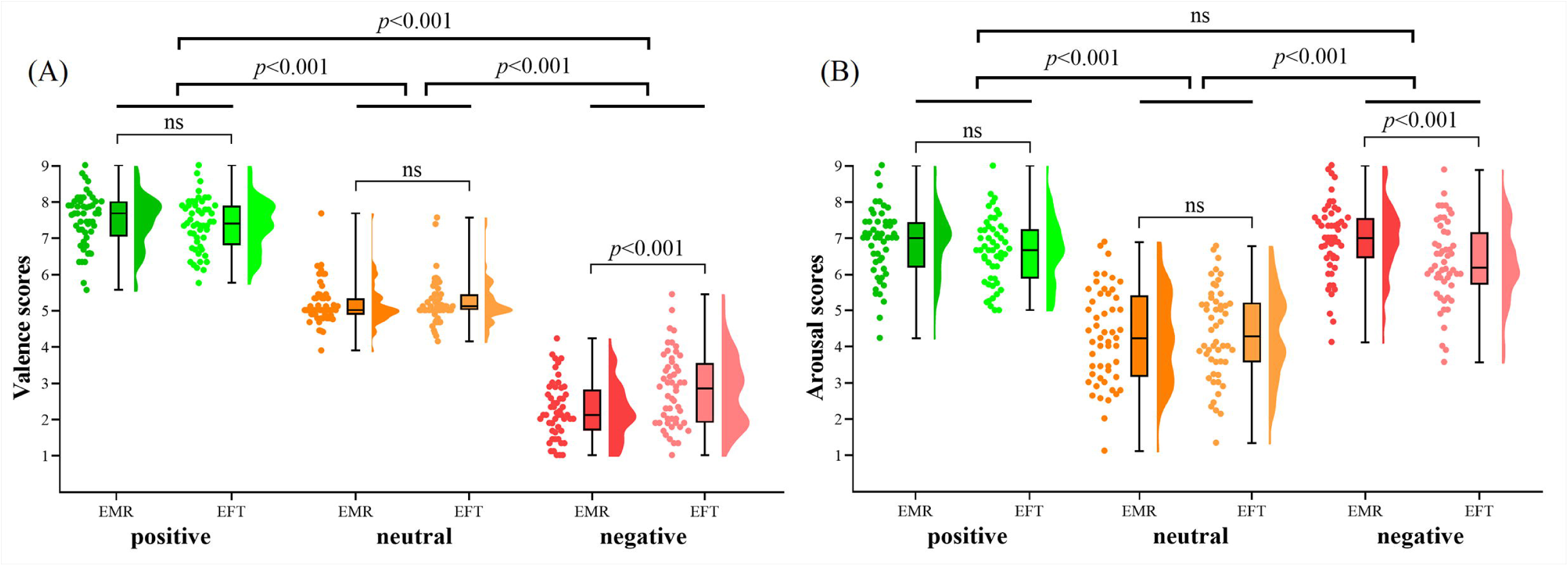
Behavioural results. Rating scores of valence (**A)** and arousal (**B**) during the tasks of episodic memory recall (EMR) and episodic future thinking (EFT) for the three valence conditions (positive, neutral, and negative).

Arousal results also revealed a significant main effect of valence (F(2,102)=139.080, *p*<0.001, 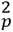=0.182) and a significant main effect of temporal context (F(1,51)=11.327, *p*=0.001, 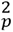=0.732), as well as a significant valence×temporal context interaction effect (F(2,102)=11.466, *p*<0.001, 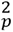=0.184; Fig 2B). Post-hoc analyses revealed significantly lower arousal ratings in the neutral relative to both positive and negative conditions across past and future scenarios (all *ps*<0.001), and higher arousal ratings in the positive than the negative were only found in the future condition (*p*=0.048), indicating a more intense feeling for positive events relative to negative conditions in the future scenarios. Significantly higher arousal ratings in the past relative to the future were only found in the negative condition (*p*<0.001), which may indicate past negative events evoking a stronger emotional response than the future scenarios.

For vividness ratings, we found a significant main effect of valence (F(2,102)=30.887, *p*<0.001, 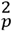=0.377) and temporal context (F(1,51)=59.314, *p*<0.001, 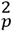=0.538), and also found a significant interaction effect of valence×temporal context (F(2,102)=7.207, *p*=0.001, 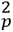=0.124). Post-hoc analyses revealed higher vividness ratings both in the positive and negative conditions relative to the neutral ones across past and future scenarios (all *ps*<0.002). Significantly higher vividness ratings in the past compared to the future across the positive, neutral, and negative conditions (all *ps*<0.026, see S1 Table).

### 2.2 Whole-brain analysis

A flexible factorial ANOVA was firstly conducted to examine the valence-specific and common brain activation patterns for the temporal contexts revealed that during past positive (relative to neutral) memory retrieval, the vmPFC, nucleus accumbens (NAcc), PCC (extending to the precuneus), anterior cingulate cortex (ACC), precentral gyrus and temporal and frontal regions were involved in memory retrieval (see Fig 3A and S2 Table; *p*_FDR_<0.05). Recalling past negative (relative to neutral) events revealed activity in a network encompassing dorsomedial prefrontal cortex (dmPFC; extending from the ACC), dlPFC, and other temporal and frontal regions engaged in memory retrieval (see Fig 3B and S3 Table). While examining activation for future imagination of positive (relative to neutral) events revealed no activation on the correction of FDR, survived in the vmPFC (extending to the ACC), PCC (extending to the precuneus), and other frontal and temporal regions involved in episodic foresight (*p*_uncorrected_<0.005; Fig 3C and S4 Table). For ‘negative>neutral’ in the future scenarios, significant activation in the dmPFC (extending to right dlPFC), PCC/precuneus, precentral gyrus, and temporal and frontal regions was found (see Fig 3D and S5 Table). Moreover, we further showed a river plot to illustrate the spatial similarity between these contrasts (*p*_uncorrected_<0.005) and a set of a priori regions of interest that have been significantly involved in mental time travel in previous studies (for detailed information on the ROIs, refer to S6 Table; see also [1,33,34]). As depicted in Fig 3E, the mPFC and PCC predominantly contribute to each contrast, whereas the ventral portion was primarily activated by positive mental imagery, and the dorsal part was engaged in negative affective events across both the EMR and EFT tasks.

**Fig 3.**
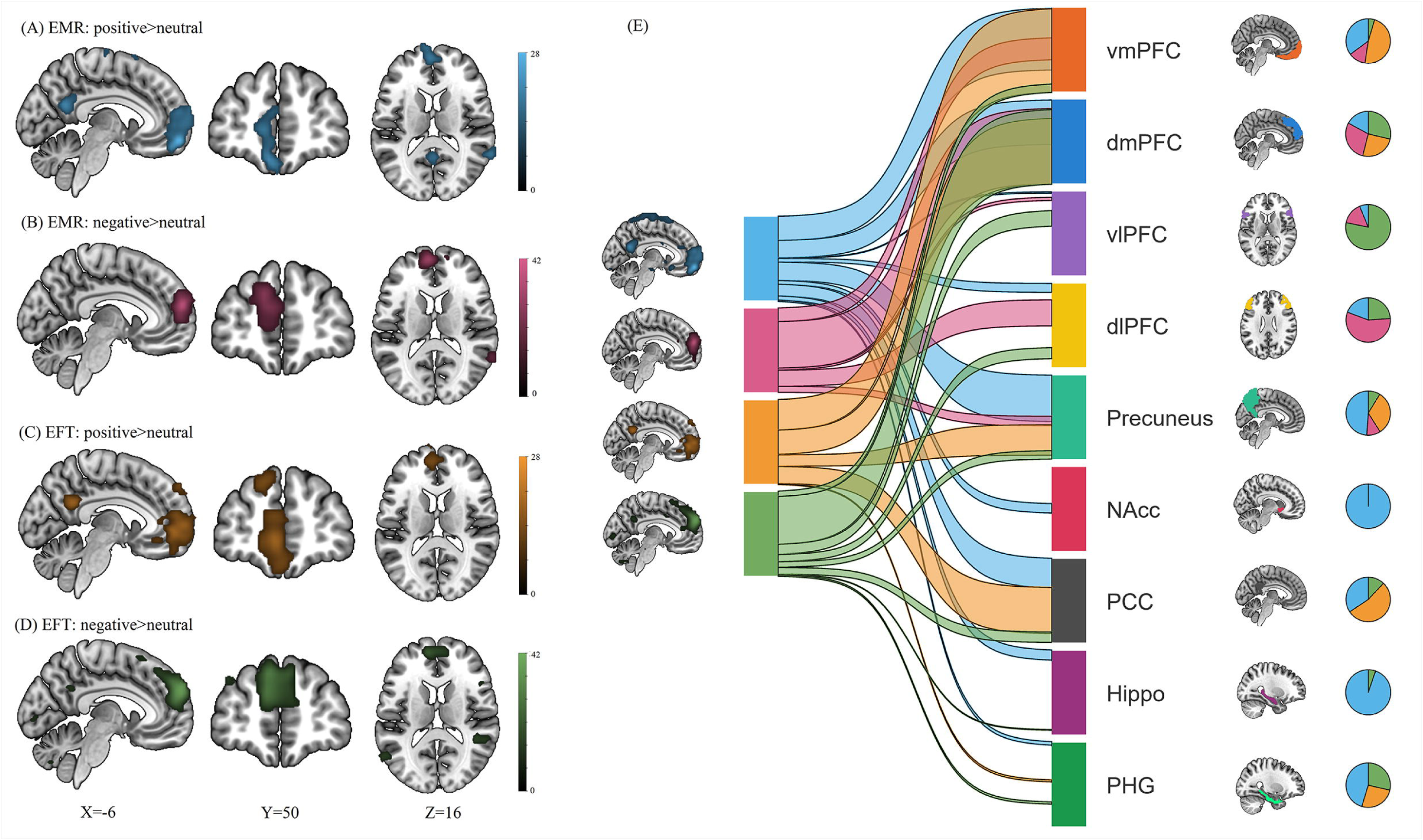
Neural activity during episodic memory recall (EMR) or episodic future thinking (EFT) according to valence. **A** Whole-brain maps of regions exhibiting significantly greater activity during positive relative to neutral in the EMR and in the EFT (**C**). **B** Whole-brain maps of regions exhibiting significantly greater activity during negative relative to neutral in the EMR and in the EFT (**D**). Note that statistical thresholds were set at *p*_FDR_<0.05 for A-D. **E** River plot illustrating the spatial similarity of neural patterns across the four contrasts (*p*_uncorrected_<0.005).

Temporal context specificity was further examined on contrast ‘EMR>EFT’, showing stronger activity in the PCC/precuneus, and superior occipital gyrus during the EMR task compared to the EFT task (S7 Table; *p*_FDR_<0.05).

In addition, one-way ANOVAs were conducted across all valence conditions for the EMR and EFT tasks, respectively, revealing the vmPFC (extending to the ACC and dmPFC), PCC, precentral gyrus, precuneus, and temporal and frontal regions involved in the EMR task, the dmPFC (extending to right dlPFC), PCC, precentral gyrus, and other middle temporal regions engaged in the EFT task. A conjunction analysis was further conducted to identify regions that contributed in both temporal context conditions, revealing activation in the dmPFC, MTG, and PCC/precuneus (*p*_FDR_<0.05; see S8 Table).

### 2.3 Parametric analysis

To identify brain regions modulated by arousal levels, parametric analysis was conducted and revealed that activation in the dlPFC, vlPFC, PCC, hippocampus, putamen, MCC (extending to the dorsal anterior cingulate cortex), thalamus, superior frontal gyrus, anterior insula, and middle frontal gyrus (MFG) increased with arousal, while no region was found in decreased activation with increasing arousal (*p*_FWE_<0.001; see Fig 4A and S9 Table).

**Fig 4.**
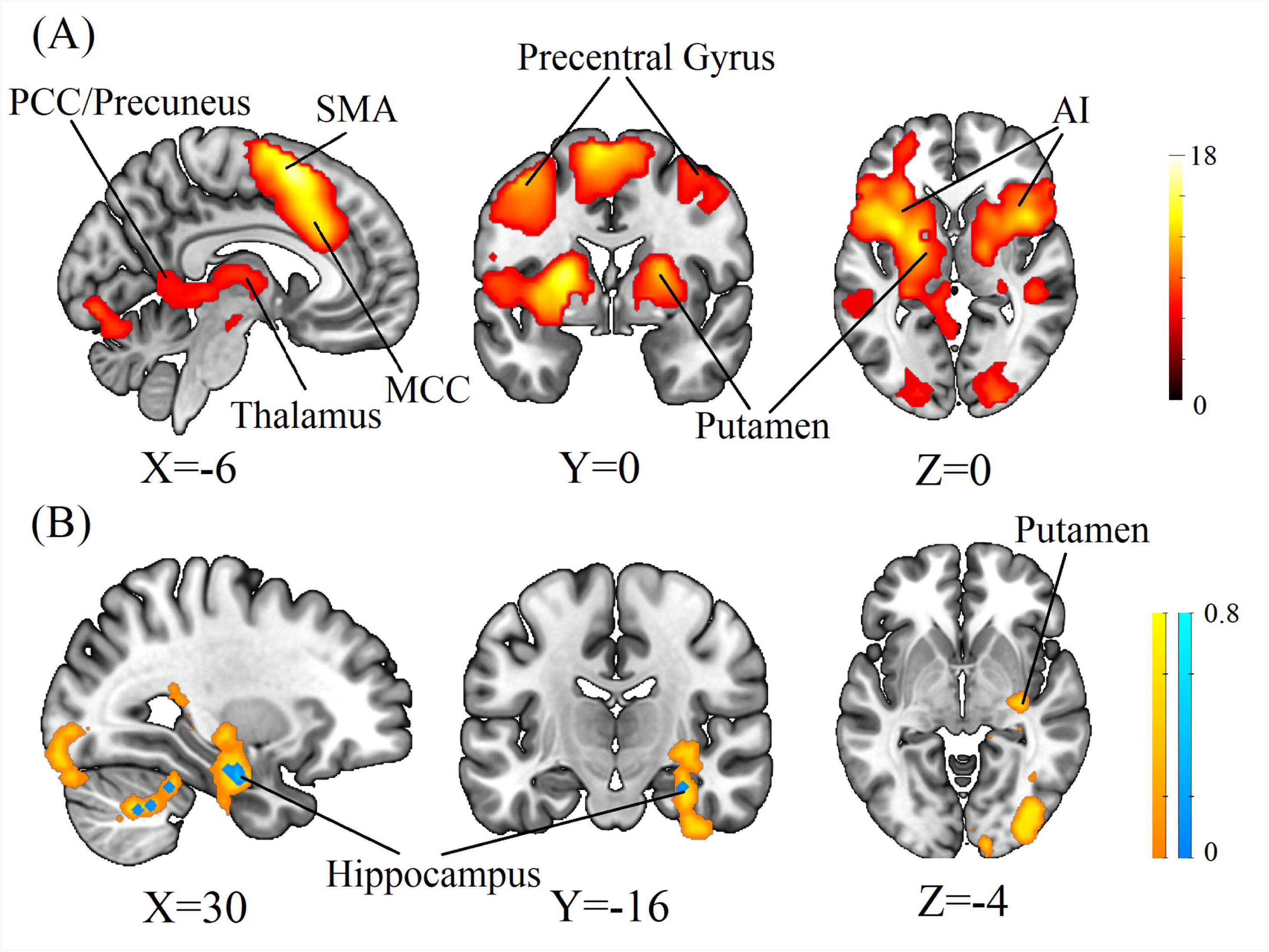
Neural results of parametric analysis and a voxel-wise searchlight representational similarity analysis. **A** Results of the weighted parametric analysis using arousal rating scores (*p*_FWE_<0.001). **B** A voxel-wise searchlight representational similarity analysis using representational dissimilarity matrix between behavioural rating (yellow color indicates valence and blue color indicates arousal) and univariate neural activity for six separate conditions showing regions whose activity patterns had a significantly positive correlation with valence scores and arousal scores (*p*_FDR_<0.001). AI: anterior insula. MCC: middle cingulate cortex. PCC: posterior cingulate cortex. SMA: supplementary motor cortex.

### 2.4 Functional connectivity analysis

To explore shared changes of functional connectivity (FC) between target ROIs and other brain regions during past and future scenarios, we conducted a conjunction analysis across all valence conditions for the EMR and EFT tasks, respectively. For the vmPFC as the target ROI, results showed strengthened FC between the vmPFC and core regions of the DMN and memory networks, including the PCC/precuneus, hippocampus, parahippocampal gyrus, and angular gyrus, across all valence conditions in the EMR task (*p*_FDR_<0.05). During the EFT task, significant increases in FC enhancement emerged between the vmPFC and the dmPFC, PCC/precuneus, MFG/precentral gyrus, and angular gyrus. For the PCC as the target ROI, results showed increased FC with the precentral gyrus, middle temporal gyrus and inferior frontal gyrus during the EMR task (*p*_FDR_<0.05), and a trend-level PCC/precuneus and angular gyrus coupling (*p*_uncorrected_<0.005) that did not survive FDR correction during the EFT task.

### 2.5 Representational similarity analysis

The searchlight-based RSA using RDM from neural data and valence or arousal ratings was conducted, revealing the right hippocampus, putamen, fusiform, and inferior occipital gyrus associated with valence differences, and also the right hippocampus associated with arousal differences for 6 conditions (see Fig 4B).

## Discussion

The present study employed an episodic memory and episodic foresight paradigm to explore the shared and distinct neurocognitive brain correlates underpinning affective EMR and EFT processes. Converging evidence from univariate, functional connectivity, and representational similarity analyses reveals a core neural pattern representing mental time travel, encompassing a common and valence-invariant engagement of core DMN regions, including the mPFC, PCC, and MTG. Notably, these regions underpin self-referential construction across past experiences or future events, supporting a generalizable system for autobiographical processes. In addition, our findings indicate that distinct neural patterns differentiate EMR and EFT processes, while affect modulates recruitment of medial frontal systems, with positive valence processes engaging more ventromedial and negative emotional processes recruiting more dorsomedial systems. The findings demonstrate that common and specific brain systems underlie mental time travel and the affective content of the time travel recruits distinguishable medial prefrontal systems.

In line with previous findings, our results also demonstrate a similar behavior and neural pattern across EMR and EFT [2,14]. At the behavioral level, participants’ subjective pleasure (i.e. positive>neutral>negative) and arousal (i.e. positive/negative>neutral) were modulated by valence across the temporal context. At the neural level, the results of the whole-brain analyses showed that different valence conditions in the EMR or EFT task compared to the neutral condition mainly activated the mPFC, PCC, and MTL. Our conjunction analysis across temporal contexts further revealed common activation in the dmPFC, PCC/precuneus, and MTG, and as such confirmed our primary hypothesis suggesting that shared recruitment of DMN core regions will contribute to EMR and EFT across valence conditions. The engagement of these cortical midline DMN hubs, irrespective of temporal direction or emotional content, may further emphasize the fundamental role of DMN for self-referential processing. The mPFC and PCC extending into the adjacent precuneus play crucial roles in integrating self-relevant information [35,36] and support episodic memory and simulation [37]. Critically, it may indicate that this core network across all valence conditions provides a basic but general cognitive content across emotional contents, and its interactions with other affective and sensory regions contribute to emotional simulation of EFT together [38,39].

Our findings examining neural recruitment across specific temporal contexts revealed stronger recruitment of the PCC extending to the precuneus and superior occipital gyrus for EMR relative to EFT, supporting our second hypothesis proposing that distinct neural systems contribute to remembering and future simulation. Consistent with the sensory reinstatement hypothesis, recalling past events involves the reactivation of sensory-perceptual details stored in posterior cortical regions [40,41]. The heightened visual cortical activity during the EMR likely reflects the greater reliance on stored sensory engrams. Conversely, while the univariate contrast of EFT compared to EMR did not show robust results after FDR-correction in frontoparietal regions, our functional connectivity findings provide support for the hypothesis that EFT would also lead to a stronger engagement of frontoparietal regions, possibly reflecting the functional coupling between the DMN and frontoparietal regions that are collaboratively supporting this cognitive process.

The neural substrates of emotion processing were distinctly different between past and future events. Recalling positive memories robustly engaged key regions of reward processing and valuation, including the vmPFC and NAcc. Similar to positive memory retrieval, the neural systems underlying future thinking of positive events also revealed activation in the vmPFC which may relate to the role of this region in positive valence encoding [21,42,43], although these results did not survive robust statistical correction, suggesting a more individually variable or distinct neural signature for anticipated versus remembered pleasure.

The difference observed across temporal contexts may have clinical implications for emotional disorders, which are characterized by dysfunction of the ability to recall pleasant experiences or generate vivid and positive future simulations, such as in depressive disorders. Thus, targeting the vmPFC using various interventions, such as transcranial magnetic stimulation (e.g., [44,45]) or real-time fMRI neurofeedback training (e.g., [46,47]), to elicit its strong activation may be a promising way to improve negative biases across temporal dimensions and in turn depressive symptoms. Memory retrieval for negative events engaged regions associated with emotional regulation and cognitive control, including the dmPFC [21,48] and dlPFC [49], possibly reflecting the effort required to process and contextualize negative autobiographical memories. Future negative simulations, however, engaged the posterior insula that is critical for interoceptive awareness and negative feeling [50,51], suggesting its role in sensory processing of negative stimuli. Consequently, these findings may suggest the need for individualized intervention strategies based on the temporal focus for clinical translation. For instance, the core target for disorders characterized by intrusive memories of past trauma could be to strengthen prefrontal cognitive control (e.g., dmPFC/dlPFC) [52], whereas a potential way to regulate symptoms dominated by anticipatory anxiety about the future might focus on modulating hyperactivation of the posterior insula to threat [6,53]. Notably, our RSA results further revealed that the hippocampus and striatum (putamen) encode representationally similar aspects of valence and arousal. It may indicate that these regions do not merely linearly respond to emotional intensity, as suggested by the parametric modulation analysis showing arousal-related activation, but contain fine-grained patterns that can distinguish the unique subjective emotional experience for each scenario across emotional contents and temporal contexts [21,54–56]. These findings may underscore how the brain encodes memories and simulations with their specific affective content that is vital for generating corresponding strategies for emotion regulation and decision-making in the future. Thus, the potential clinical translation of the present study for psychopathology is to intervene in these specific circuits to alter maladaptive in mental time travel, which may induce the over-generalization and negative bias in emotional disorders (e.g., overgeneral autobiographical memory in depression [57]), including (1) hyperactivity in negative-memory networks (dmPFC/dlPFC), (2) hypoactivity in reward networks (vmPFC/NAcc) during future simulation, or (3) aberrant hippocampal-striatal coupling that over-represents negative outcomes.

There are several limitations in the present study. First, while we focused on valence and arousal, other dimensions like detail and personal significance influence mental time travel and should be explored in future studies. Second, future research could examine the generalizability and specificity of this model in clinical populations since our sample only consisted of healthy young adults. Future studies could utilize longitudinal experimental designs to track how these neural patterns alter with therapeutic or aging interventions.

In summary, the present study delineated both a common core and dissociable neural systems by decomposing mental time travel into its temporal and affective components. Our findings confirmed the central role of the DMN in self-projection, highlighted a dissociation between valence and temporal context, and revealed how the brain encodes the unique emotions of our past and future selves through multivariate patterning. Moreover, our findings underscore that across the time domain the medial prefrontal cortex encodes the valence of the content, with ventral regions encoding positive and more dorsal regions encoding negative emotional experience. The present findings provide a robust empirical insight for understanding the human capacity to mentally experience time and reveal new neural targets for alleviating mental disorders characterized by dysfunction of these psychological processes.

## Materials and Methods

### 4.1 Participants

62 healthy right-handed participants (31 males, mean age=21.74 years, SD=2.26) from the University of Electronic Science and Technology of China (UESTC) were enrolled in the present study. Given that the neural activity during internal mental processes critically relies on the immersive participation of the volunteers we initially performed a quality assessment examining performance in the memory recall or foresight task (n=9), one participant was additionally excluded due to not completing the whole experiment (n=1), leading to a final sample size of 52 participants in data analyses (26 males, mean age=21.56 years, SD=2.13). All participants gave informed consent and reported no history of or current psychiatric or neurological disorders before MRI scanning. All procedures of the current study were approved by the local ethical committee at UESTC and adhered to the latest version of the Declaration of Helsinki. This study was preregistered on the Open Science Framework (https://osf.io/v3kdc).

### 4.2 Experimental procedure

The experimental protocol comprised two visits for all participants (see Fig 1A). On the first visit, participants were instructed to complete an autobiographical interview task (details see below) and a series of validated scales to assess anxiety and depression levels, including the Beck Depression Inventory-II (BDI-II; [58]) and the State-Trait Anxiety Inventory (STAI; [59]). In addition, all participants completed the Positive and Negative Affect Schedule (PANAS; [60]) before and after MRI scanning to assess mood changes induced by the recall paradigm.

On the second visit participants were asked to perform two memory test tasks (details see S1 Text) to determine whether they clearly remembered the corresponding events. All participants expressed that they understood the experimental procedure and also performed a practice task to familiarize the EMR and EFT tasks before receiving MRI scanning room.

#### 4.2.1 Autobiographical Interview Task

In line with previous studies [16,61], participants were provided with a list of life event cues to recall or imagine 5 affective events involved in strong personal emotional experience for each valence condition spanning past and future scenarios. Specifically, they were instructed to recall the 5 most positive (e.g., joyful or exciting), 5 neutral (e.g., daily or calm), and 5 most negative (e.g., sad or angry) events from the past 5 years, and also to imagine the 5 most positive, 5 neutral, and 5 most negative events that might happen in the future 5 years. The list served as a guide to facilitate recall or imagination, but participants were also free to recall or imagine events not included in the list. They were instructed to write 3-5 sentences describing concrete details (e.g., what, when, where, and with whom) for each affective event within the past or future 5 years. Subsequently, 3 memory cue words were generated for each event to facilitate a specific and vivid recall/imagination during the second visit. After writing materials, their rating scores of valence (How did you feel when you recall/imagine this event? 1=very negative, 5=neutral, 9=very positive), arousal (How emotionally intense when you recall/imagine this event? 1=not at all intense, 9=very intense), and vividness (How vividly for this event when you recall/imagine it? 1=not at all vivid, 9=very vivid) were also record on a 9-point Likert scale. Based on participants’ subjective ratings, the 3 events for each condition were included to ascertain the affective validity of the experimental materials during MRI scanning. In total, 18 events were included in the second visit. The cue words of each event were presented to assist participants in engaging in EMR and EFT tasks.

#### 4.2.2 Episodic Memory Retrieval (EMR) and Episodic Future Thinking (EFT) Tasks

After an anatomical scan and resting-state fMRI scan, all participants performed EMR and EFT tasks. Each task comprised 3 runs, with EMR and EFT runs displayed in alternation. Each run included 9 affective events, with 3 events per valence condition. The order of EMR and EFT tasks, as well as the valence order, was counterbalanced across all participants. The experimental procedure of EMR or EFT tasks is shown in Fig 1B. After a jittered fixation cross of 8-12 s, they were asked to recall or imagine the personal scenarios based on the 3 cue words (e.g., family, vocation, seaside) from the first visit. Within a time window of 16 s, participants indicated the duration of the memory by button press from their first button press indicated the ‘start’ (i.e. they started to engage in EMR or EFT), to the second button press indicated the ‘end’ (i.e. they finished this trial of EMR or EFT). After a jittered fixation of 1-2 s, participants were instructed to indicate whether they could vividly recall the event within 2 s (Yes/No), with additional questions on the emotional experience following after 1-2 s to assess their rating scores of valence and arousal for each event on a 9-point Likert scale within a duration of 4 s. The present paradigm included emotional valence (three levels: positive, neutral, and negative) and temporal context (two levels: EMR and EFT) as within-subject factors. After MRI acquisition, participants performed a vividness assessment on a 9-point Likert scale for each affective event presented in the EMR and EFT tasks.

### 4.3 Image data acquisition

Images were collected using a 3T, GE Discovery MR750 system (General Electric Medical System, Milwaukee, WI, USA). High-resolution whole-brain volume T1-weighted images were collected with a 3D spoiled gradient echo pulse sequence (repetition time, 6 ms; echo time, 2 ms; flip angle, 12°; field of view=256 × 256 mm; acquisition matrix, 256 × 256; thickness, 1 mm; number of excitations, 2; 156 slices). Functional images were acquired using a T2-weighted echo-planar imaging pulse sequence (repetition time, 2000 ms; echo time, 30 ms; slices, 39; thickness, 3 mm; gap, 1 mm; field of view, 240 × 240 mm; resolution, 64 × 64; flip angle, 90°).

### 4.4 Data analyses

#### 4.4.1 Imaging Data Analyses

SPM12 (Wellcome Department of Cognitive Neurology, London, UK, http://www.fil.ion.ucl.ac.uk/spm) was used for image processing. The first 5 volumes were removed to allow magnet steady states. The remaining images were preprocessed, including head-motion correction, co-registration of the mean functional image and the T1 image, normalization to the standard Montreal Neurological Institute (MNI) space, and smoothing using an 8 mm full-width at half maximum (FWHM). For the EMR and EFT tasks, the first-level design matrix included 9 regressors (positive, neutral, negative, button press of ‘start’, button press of ‘end’, determination choice, valence rating, arousal rating, and invalid trials) convolved with the canonical hemodynamic response function (HRF) and the 6 head motion parameters. In the invalid condition, trials were excluded due to reasons such as participants selecting “no” during the determination choice phase or trials with a duration shorter than 2s in the EMR or EFT tasks. Contrast images were generated for each experimental condition (‘positive EMR’, ‘neutral EMR’, ‘negative EMR’, ‘positive EFT’, ‘neutral EFT’, and ‘negative EFT’), as well as for valence comparisons (i.e. ‘EMR: positive>neutral’, ‘EMR: negative>neutral’, ‘EFT: positive>neutral’, and ‘EFT: negative>neutral’) and temporal comparisons (i.e. EMR>EFT) for each session of the EMR and EFT tasks.

For the second level the contrasts aimed at determining the common and distinct valence- and temporally-specific neural activation patterns using appropriate first-level contrasts and conjunction analyses. Valence-specific and temporal context effects were examined based on a flexible factorial ANOVA model using a first-level contrast image of 6 separate conditions. In addition, conjunction analyses were conducted to identify the shared neural activity across valence in different temporal contexts using one-way ANOVA for all valence conditions in the past or future scenarios, respectively. At the whole-brain level, results were corrected using the False Discovery Rate peak-level correction (*p*_FDR_<0.05) and only clusters exceeding 10 voxels were reported.

#### 4.4.2 Examination of arousal

A subject-level GLM was conducted to identify regions modulated by EMR/EFT arousal ratings. The first level design matrix only included 7 regressors for the EMR or EFT tasks (retrieval/imagination trials, button press of ‘start’, button press of ‘end’, determination choice, valence rating, arousal rating, and invalid trials) convolved with the HRF and the 6 head motion parameters. Additionally, exploratory analyses were conducted using parametric modulation at the singlelJsubject level to examine the potential influence of affective arousal on the neural activation (e.g., Zhang et al. [25]). TriallJwise arousal ratings (categorized as low: 1–3, medium: 4–6, and high: 7–9) were included as parametric modulators for both the retrieval and imagination periods. A one-sample T-test was performed on contrast images for all participants. At the whole-brain level, results were corrected using the False Discovery Rate peak-level correction (*p*_FDR_<0.05). Only clusters exceeding 10 voxels were reported.

#### 4.4.3 Functional Connectivity Analyses

A psychophysiological interaction analysis was conducted using the gPPI toolbox [62] to assess changes in functional connectivity between core regions and other brain regions for different valences in EMR and EFT tasks (e.g., positive EMR). The Regions of interest (ROIs) were defined using the Yeo17 atlas [63], including the ventromedial prefrontal cortex (vmPFC) and PCC which are core components of the DMN [64,65] and frequently co-activated in the EMR and EFT processes [1,2]. Additionally, contrast images were generated using one-sample T-tests for all valence conditions in the past or future scenarios, respectively. Subsequently, conjunction analyses were conducted to identify shared neural networks associated with pre-defined ROIs across valence conditions in different temporal contexts.

#### 4.4.4 Representational Similarity Analyses

Representational similarity analysis (RSA) is a powerful method to explore the relationship between different representations, such as across modalities (e.g., between behavioral and other modalities) or across species (e.g., between humans and animals) [66,67]. An fMRI-based representation dissimilarity matrix (RDM) was conducted using a searchlight analysis to assess neural pattern similarity across 6 separate conditions using the NeuroRA toolbox in Python [68]. Additionally, two behavioral RDMs (i.e., valence RDM and arousal RDM) were calculated based on mean scores for each condition per participant. Subsequently, a searchlight-based RSA was conducted to identify brain regions closely associated with individual valence and arousal scores.

#### 4.4.5 Statistical Analyses

Two-way repeated-measures ANOVAs were conducted to assess the effects of temporal context (EMR vs. EFT) and valence (positive vs. neutral vs. negative) on subjective valence, arousal, and vividness scores. A paired-sample t-test was performed to evaluate mood effect using PANAS scores before and after MRI scanning.

## Acknowledgements

This work is supported by the National Natural Science Foundation of China (NSFC 82271583), the Ministry of Science and Technology of China (STI 2030–Major Projects 2022ZD0208500), the Hong Kong University Grants Council (GRF 17615525), the University of Hong Kong seed funding and start-up schemes (2407102536).

## Supporting information

**S1 Table. Behavioural results.** The statistical results of simple effects analyses for a mixed ANOVA on subjective rating scores of valence, arousal, and vividness. Note that EMR indicates episodic memory recall and EFT indicates episodic future thinking.

**S2 Table. Neural activity in the episodic memory retrieval task (positive vs. neutral).** Brain regions showing stronger activity in positive relative to neutral conditions in the episodic memory retrieval (EMR) (MNI coordinates). Note that all regions are reported with a *p*_FDR_<0.05 threshold at the whole-brain level. L indicates left; R indicates right.

**S3 Table. Neural activity in the episodic memory retrieval task (negative vs. neutral).** Brain regions showing stronger activity in negative relative to neutral conditions in the episodic memory retrieval (EMR) (MNI coordinates). Note that all regions are reported with a *p*_FDR_<0.05 threshold at the whole-brain level. L indicates left; R indicates right.

**S4 Table. Neural activity in the episodic future thinking task (positive vs. neutral).** Brain regions showing stronger activity in positive relative to neutral conditions in the episodic future thinking (EFT) (MNI coordinates). Note that all regions are reported with a *p*_uncorrected_<0.005 threshold at the whole-brain level. L indicates left; R indicates right.

**S5 Table. Neural activity in the episodic future thinking task (negative vs. neutral).** Brain regions showing stronger activity in positive relative to neutral conditions in the episodic future thinking (EFT) (MNI coordinates). Note that all regions are reported with a *p*_FDR_<0.05 threshold at the whole-brain level. L indicates left; R indicates right.

**S6 Table. Selected ROI information in the riverplot.** Selected ROIs for Fig. 3E.

**S7 Table. Neural activity in the episodic memory retrieval relative to the episodic future thinking task.** Brain regions showing stronger activity during episodic memory retrieval (EMR) in relative to episodic future thinking (EFT) across all valences (MNI coordinates). Note that all regions are reported with a *p*_FDR_<0.05 threshold at the whole-brain level. L indicates left; R indicates right.

**S8 Table. Neural activity in the episodic memory retrieval or episodic future thinking task.** Brain regions showing stronger activity across all valence conditions in the episodic memory retrieval (EMR) and episodic future thinking (EFT) (MNI coordinates). Note that all regions are reported with a *p*_FDR_<0.05 threshold at the whole-brain level. L indicates left; R indicates right.

**S9 Table. Brain region in parametric modulation analysis.** Brain regions showing stronger activity with higher arousal level across the episodic memory retrieval (EMR) and episodic future thinking (EFT) tasks (MNI coordinates). Note that all regions are reported with a *p*_FWE_<0.001 threshold at the whole-brain level. L indicates left; R indicates right.

