## Supplemental Materials for "Shared and distinct default mode systems in mental time travel and affective experience over time"

**S1 Text.**

**Memory test tasks**

All participants were asked to perform two memory test tasks to determine the availability of cue words they wrote, including (1) a memory recognition task and (2) a memory recall task. During the memory recognition task with 18 trials, a cue word was displayed at first in a trial, then they were asked to choose the correct one from 4 words, including another cue word from this event and 3 cue words unrelated to this event. During the memory recall task, they were required to retell two affective event contents after the experimenter showed one or two cue words from 2 events that were not included in the later MRI scanning. The correct rate is over 80% for all participants who were recruited in the present study.

**S1 Table.** **Behavioural results.** The statistical results of simple effects analyses for a mixed ANOVA on subjective rating scores of valence, arousal, and vividness.

| Comparison | Mean Difference | Standard Error | p-value |
| --- | --- | --- | --- |
| **Valence** |  |  |  |
| EMR: positive vs. neutral | 2.321 | 0.118 | <0.001 |
| EMR: positive vs. negative | 5.241 | 0.194 | <0.001 |
| EMR: neutral vs. negative | 2.921 | 0.143 | <0.001 |
| EFT: positive vs. neutral | 2.156 | 0.118 | <0.001 |
| EFT: positive vs. negative | 4.568 | 0.203 | <0.001 |
| EFT: neutral vs. negative | 2.412 | 0.179 | <0.001 |
| Positive: EMR vs. EFT | 0.122 | 0.079 | 0.127 |
| Neutral: EMR vs. EFT | -0.042 | 0.077 | 0.582 |
| Negative: EMR vs. EFT | -0.551 | 0.135 | <0.001 |
| **Arousal** |  |  |  |
| EMR: positive vs. neutral | 2.600 | 0.204 | <0.001 |
| EMR: positive vs. negative | -0.090 | 0.112 | 1.000 |
| EMR: neutral vs. negative | -2.690 | 0.222 | <0.001 |
| EFT: positive vs. neutral | 2.310 | 0.199 | <0.001 |
| EFT: positive vs. negative | 0.346 | 0.139 | 0.048 |
| EFT: neutral vs. negative | -1.963 | 0.193 | <0.001 |
| Positive: EMR vs. EFT | 0.208 | 0.108 | 0.059 |
| Neutral: EMR vs. EFT | -0.082 | 0.108 | 0.452 |
| Negative: EMR vs. EFT | 0.645 | 0.133 | <0.001 |
| **Vividness** |  |  |  |
| EMR: positive vs. neutral | 1.561 | 0.196 | <0.001 |
| EMR: positive vs. negative | 0.196 | 0.159 | 0.674 |
| EMR: neutral vs. negative | -1.366 | 0.218 | <0.001 |
| EFT: positive vs. neutral | 1.073 | 0.218 | <0.001 |
| EFT: positive vs. negative | 0.416 | 0.218 | 0.187 |
| EFT: neutral vs. negative | -0.657 | 0.182 | 0.002 |
| Positive: EMR vs. EFT | 0.812 | 0.147 | <0.001 |
| Neutral: EMR vs. EFT | 0.323 | 0.141 | 0.026 |
| Negative: EMR vs. EFT | 1.032 | 0.146 | <0.001 |

Note that EMR indicates episodic memory recall and EFT indicates episodic future thinking.

**S2 Table.** **Neural activity in the episodic memory retrieval task** **(positive vs. neutral)**.

Brain regions showing stronger activity in positive relative to neutral conditions in the episodic memory retrieval (EMR) (MNI coordinates).

| Brain Region | BA | No. Voxels | Peak F-value | X | Y | Z |
| --- | --- | --- | --- | --- | --- | --- |
| L. Ventromedial Prefrontal Cortex | 10/11/32 | 427 | 27.40 | -6 | 59 | -13 |
| Medial Frontal Gyrus |  |  | 20.85 | -9 | 65 | 8 |
| Medial Frontal Gyrus |  |  | 19.08 | -3 | 56 | 14 |
| Anterior Cingulate Cortex |  |  | 18.59 | -9 | 50 | 2 |
| L. Inferior Temporal Gyrus | 20/21 | 76 | 24.06 | -60 | -10 | -19 |
| Inferior Temporal Gyrus |  |  | 16.76 | -63 | -4 | -28 |
| L. Posterior Cingulate Cortex | 31 | 164 | 23.70 | -3 | -58 | 23 |
| R. Middle Temporal Gyrus |  | 79 | 22.74 | 63 | -1 | -28 |
| R. Superior Temporal Gyrus |  | 38 | 17.66 | 63 | -49 | 14 |
| L. Superior Frontal Gyrus | 6 | 84 | 16.90 | -12 | -16 | 71 |
| Precentral Gyrus |  |  | 16.40 | -24 | -16 | 74 |
| L. Superior Frontal Gyrus |  | 18 | 16.38 | -24 | 38 | 47 |
| R. Inferior Temporal Gyrus |  | 24 | 16.34 | 60 | -70 | -1 |
| L. Nucleus Accumbens |  | 11 | 14.58 | -24 | 11 | -16 |
| L. Superior Frontal Gyrus |  | 20 | 14.43 | -9 | 17 | 68 |
| Supplementary Motor Area |  |  | 12.73 | -3 | 8 | 74 |
| L. Middle Temporal Gyrus |  | 17 | 14.27 | -48 | -67 | 26 |
| Precuneus |  | 22 | 13.60 | 0 | -52 | 65 |

Note that all regions are reported with a *p*_FDR_<0.05 threshold at the whole-brain level. L indicates left; R indicates right.

**S3 Table.** **Neural activity in the episodic memory retrieval task** **(negative vs. neutral).**

Brain regions showing stronger activity in negative relative to neutral conditions in the episodic memory retrieval (EMR) (MNI coordinates).

| Brain Region | BA | No. Voxels | Peak F-value | X | Y | Z |
| --- | --- | --- | --- | --- | --- | --- |
| L. Dorsomedial Prefrontal Cortex | 9/10 | 455 | 36.22 | -9 | 56 | 20 |
| Dorsomedial Prefrontal Cortex |  |  | 23.96 | 12 | 56 | 23 |
| L. Middle Temporal Gyrus | 21/38 | 228 | 27.99 | -54 | 5 | -34 |
| Middle Temporal Gyrus |  |  | 27.22 | -57 | 5 | -22 |
| Superior Temporal Gyrus |  |  | 24.76 | -42 | 23 | -25 |
| R. Dorsolateral Prefrontal Cortex | 9/46 | 76 | 22.86 | 54 | 35 | 26 |
| Dorsolateral Prefrontal Cortex |  |  | 22.06 | 51 | 32 | 35 |
| R. Middle Temporal Gyrus | 21/38 | 99 | 20.66 | 51 | 8 | -31 |
| Superior Temporal Gyrus |  |  | 19.97 | 57 | 11 | -25 |
| Superior Temporal Gyrus |  |  | 18.58 | 48 | 20 | -28 |
| R. Superior Temporal Gyrus | 22 | 36 | 18.73 | 63 | -49 | 11 |
| R. Angular Gyrus | 39 | 18 | 16.11 | 45 | -79 | 29 |
| R. Middle Temporal Gyrus | 37 | 23 | 15.39 | 54 | -64 | 2 |

Note that all regions are reported with a *p*_FDR_<0.05 threshold at the whole-brain level. L indicates left; R indicates right.

**S4 Table.** **Neural activity in the episodic future thinking task (positive vs. neutral).**

Brain regions showing stronger activity in positive relative to neutral conditions in the episodic future thinking (EFT) (MNI coordinates).

| Brain Region | BA | No. Voxels | Peak F-value | X | Y | Z |
| --- | --- | --- | --- | --- | --- | --- |
| L. Posterior Cingulate Cortex | 31 | 95 | 17.90 | -3 | -49 | 26 |
| Precuneus |  |  | 14.99 | -3 | -55 | 26 |
| L. Ventromedial Prefrontal Cortex | 10/11 | 623 | 17.79 | -6 | 50 | -4 |
| Superior Frontal Gyrus |  |  | 17.64 | -9 | 59 | 2 |
| Ventromedial Prefrontal Cortex |  |  | 13.86 | -9 | 35 | -10 |
| R. Middle Temporal Gyrus | 21 | 34 | 15.79 | 42 | 5 | -31 |
| L. Superior Frontal Gyrus | 9 | 107 | 13.52 | -12 | 50 | 38 |
| Dorsomedial Prefrontal Cortex |  |  | 13.00 | -6 | 56 | 44 |
| L. Middle Temporal Gyrus |  | 21 | 10.47 | -33 | 8 | -34 |
| Middle Temporal Gyrus |  |  | 8.37 | -42 | 8 | -31 |

Note that all regions are reported with a *p*_uncorrected_<0.005 threshold at the whole-brain level. L indicates left; R indicates right.

**S5 Table.** **Neural activity in the episodic future thinking task (negative vs. neutral).**

| Brain Region | BA | No. Voxels | Peak F-value | X | Y | Z |
| --- | --- | --- | --- | --- | --- | --- |
| L. Dorsomedial Prefrontal Cortex | 8/9/10 | 613 | 41.34 | -9 | 59 | 26 |
| Dorsolateral Prefrontal Cortex |  |  | 21.19 | 6 | 50 | 35 |
| Medial Frontal Gyrus |  |  | 13.19 | 6 | 38 | 35 |
| R. Middle Temporal Gyrus | 21/38 | 809 | 28.09 | -57 | 5 | 47 |
| Middle Temporal Gyrus |  |  | 23.16 | -48 | -28 | -19 |
| Middle Temporal Gyrus |  |  | 22.12 | -45 | 5 | -7 |
| R. Superior Temporal Gyrus | 21/38 | 337 | 23.94 | 48 | 23 | -43 |
| Middle Temporal Gyrus |  |  | 21.11 | 54 | -7 | -19 |
| Inferior Frontal Gyrus |  |  | 19.10 | 27 | 20 | -16 |
| R. Superior Temporal Gyrus | 13 | 60 | 1653 | 42 | -43 | -22 |
| R. Inferior Temporal Gyrus | 21/38 | 75 | 15.57 | 42 | 8 | 11 |
| Middle Temporal Gyrus |  |  | 15.18 | 48 | 8 | -46 |
| L. Cuneus | 17 | 21 | 15.46 | -12 | -88 | -34 |
| L. Ventrolateral Prefrontal Cortex |  | 14 | 14.68 | -36 | 50 | 32 |
| R. Precentral Gyrus | 6 | 69 | 14.30 | 48 | -10 | 32 |
| Precentral Gyrus |  |  | 13.96 | 39 | -7 | 47 |
| Precentral Gyrus |  |  | 11.20 | 30 | -16 | 65 |
| L. Posterior Cingulate Cortex  /Precuneus |  | 15 | 12.87 | -6 | -52 | 32 |
| L. Superior Frontal Gyrus |  | 10 | 12.00 | -6 | 17 | 62 |
| R. Inferior Frontal Gyrus |  | 11 | 11.75 | 51 | 20 | 17 |
| R. Inferior Occipital Gyrus | 18 | 19 | 11.39 | 42 | -85 | -22 |

Brain regions showing stronger activity in negative relative to neutral conditions in the episodic future thinking (EFT) (MNI coordinates).

Note that all regions are reported with a *p*_FDR_ < 0.05 threshold at the whole-brain level. L indicates left; R indicates right.

| ROI | Areas included | Atlas |
| --- | --- | --- |
| vmPFC | 10r, 10v, 10d, 10pp, OFC | HCP-MMP v1.0, Glasser et al. 2016 [1] |
| dmPFC | 8BM, 8BL, 9m | HCP-MMP v1.0, Glasser et al. 2016 [1] |
| vlPFC | 44,45 | HCP-MMP v1.0, Glasser et al. 2016 [1] |
| dlPFC | 8Av, p9-46v, 46 | HCP-MMP v1.0, Glasser et al. 2016 [1] |
| Precuneus | - | Automated Anatomical Labeling 3 (AAL3) atlas [2] |
| NAcc | - | Automated Anatomical Labeling 3 (AAL3) atlas [2] |
| PCC | - | Automated Anatomical Labeling 3 (AAL3) atlas [2] |
| Hippocampus | - | Automated Anatomical Labeling 3 (AAL3) atlas [2] |
| Parahippocampus | - | Automated Anatomical Labeling 3 (AAL3) atlas [2] |

**S6 Table.** **Selected ROI information in the riverplot.**

Selected ROIs for Fig. 3E.

**S7 Table.** **Neural activity in the episodic memory retrieval relative to the episodic future thinking task.**

Brain regions showing stronger activity during episodic memory retrieval (EMR) in relative to episodic future thinking (EFT) across all valences (MNI coordinates).

| Brain Region | BA | No. Voxels | Peak F-value | X | Y | Z |
| --- | --- | --- | --- | --- | --- | --- |
| R. Posterior Cingulate Cortex/Precuneus |  | 33 | 27.62 | 15 | -55 | 17 |
| L. Posterior Cingulate Cortex/Precuneus |  | 32 | 26.76 | -18 | -58 | 17 |
| L. Superior Occipital Gyrus |  | 19 | 22.97 | -36 | -85 | 29 |

Note that all regions are reported with a *p*_FDR_<0.05 threshold at the whole-brain level. L indicates left; R indicates right.

**S8 Table.** **Neural activity in the episodic memory retrieval or episodic future thinking task.**

Brain regions showing stronger activity across all valence conditions in the episodic memory retrieval (EMR) and episodic future thinking (EFT) (MNI coordinates).

| Brain Region | BA | No. Voxels | Peak F-value | X | Y | Z |
| --- | --- | --- | --- | --- | --- | --- |
| **EMR: Main effect** |  |  |  |  |  |  |
| L. Dorsomedial Prefrontal Cortex | 10/11 | 588 | 22.96 | -6 | 56 | 17 |
| Ventromedial Prefrontal Cortex |  |  | 15.63 | -6 | 59 | -13 |
| Dorsolateral Prefrontal Cortex |  |  | 11.80 | 12 | 56 | 23 |
| L.Middle Temporal Gyrus | 21/38 | 270 | 17.08 | -60 | 5 | -25 |
| Inferior Temporal Gyrus |  |  | 12.24 | -60 | -10 | -19 |
| R. Superior Temporal Gyrus | 22/37 | 121 | 14.62 | 63 | -52 | 14 |
| Middle Temporal Gyrus |  |  | 10.56 | 57 | -64 | 2 |
| R.Middle Frontal Gyrus | 46 | 57 | 13.77 | 54 | 32 | 29 |
| L. Superior Frontal Gyrus | 6 | 47 | 13.21 | -15 | 17 | 65 |
| R. Tuber |  | 17 | 11.92 | 24 | -79 | -37 |
| R. Middle Temporal Gyrus | 21 | 69 | 11.44 | 63 | -1 | -31 |
| Middle Temporal Gyrus |  |  | 8.97 | 51 | 8 | -31 |
| Superior Temporal Gyrus |  |  | 8.71 | 57 | 11 | -25 |
| L. Posterior Cingulate Cortex | 23/30 | 89 | 11.22 | -3 | -55 | 23 |
| L. Superior Frontal Gyrus |  | 13 | 9.35 | -12 | -13 | 71 |
| L. Precentral Gyrus | 6 | 21 | 9.35 | -27 | -16 | 71 |
| Precuneus |  | 14 | 8.23 | 0 | -52 | 65 |
| Postcentral Gyrus |  |  | 7.51 | -3 | -43 | 71 |
| **EFT: Main effect** |  |  |  |  |  |  |
| L. Dorsomedial Prefrontal Cortex | 9/10 | 425 | 25.22 | -12 | 56 | 29 |
| Dorsolateral Prefrontal Cortex |  |  | 15.36 | 6 | 50 | 35 |
| Anterior Prefrontal Cortex |  |  | 12.10 | 6 | 56 | 23 |
| L. Middle Temporal Gyrus | 21/38 | 256 | 16.92 | -57 | 8 | -19 |
| Middle Temporal Gyrus |  |  | 14.99 | -48 | -28 | -7 |
| Superior Temporal Gyrus |  |  | 12.26 | -48 | 20 | -19 |
| L. Supramarginal Gyrus | 40 | 66 | 14.75 | -60 | -49 | 26 |
| R. Superior Temporal Gyrus | 38 | 168 | 14.71 | 48 | 23 | -22 |
| Middle Temporal Gyrus |  |  | 13.16 | 57 | -4 | -16 |
| Middle Temporal Gyrus |  |  | 11.46 | 54 | -13 | -13 |
| R. Middle Temporal Gyrus |  | 116 | 12.49 | 60 | -34 | -7 |
| Middle Temporal Gyrus |  |  | 9.28 | 45 | -25 | -7 |
| L. Posterior Cingulate Cortex | 31 | 51 | 12.18 | -6 | -49 | 29 |
| R. Precentral Gyrus |  | 32 | 11.45 | 48 | -10 | 47 |
| L. Middle Temporal Gyrus | 21/22 | 104 | 10.09 | -60 | -64 | 8 |
| Middle Temporal Gyrus |  |  | 9.87 | -60 | -43 | 2 |
| R. Middle Temporal Gyrus |  | 21 | 9.97 | 48 | 8 | -31 |
| L. Precentral Gyrus |  | 13 | 9.62 | -39 | 5 | 47 |
| R. Middle Frontal Gyrus |  | 10 | 8.88 | 39 | -4 | 65 |
| Lingual Gyrus |  | 13 | 8.52 | 0 | -91 | 2 |
| Lingual Gyrus |  |  | 8.00 | -9 | -88 | -1 |
| R. Superior Temporal Gyrus |  | 10 | 8.48 | 45 | -43 | 11 |
| R. Inferior Occipital Gyrus |  | 10 | 8.44 | 39 | -85 | -22 |
| **Conjunction analysis** |  |  |  |  |  |  |
| L. Dorsomedial Prefrontal Cortex | 9/10 | 196 | 18.74 | -9 | 56 | 23 |
| Anterior Prefrontal Cortex |  |  | 7.66 | -9 | 59 | 5 |
| L. Middle Temporal Gyrus | 21 | 23 | 13.92 | -57 | 8 | -22 |
| L. Middle Temporal Gyrus |  | 25 | 10.69 | -48 | 8 | -37 |
| L. Superior Temporal Gyrus |  | 21 | 10.44 | -45 | 23 | -22 |
| L. Posterior Cingulate Cortex |  | 11 | 9.05 | -6 | -55 | 26 |

Note that all regions are reported with a *p*_FDR_ < 0.05 threshold at the whole-brain level. L indicates left; R indicates right.

**S9 Table.** **Brain region in parametric modulation analysis.**

Brain regions showing stronger activity with higher arousal level across the episodic memory retrieval (EMR) and episodic future thinking (EFT) tasks (MNI coordinates).

| Brain Region | BA | No. Voxels | Peak t-value | X | Y | Z |
| --- | --- | --- | --- | --- | --- | --- |
| L. Superior Frontal Gyrus | 6/18/32 | 8969 | 17.21 | -3 | 11 | 56 |
| Putamen |  |  | 15.66 | -21 | -4 | 11 |
| Lingual Gyrus |  |  | 14.79 | -15 | -88 | -10 |
| Anterior Prefrontal Cortex |  |  | 12.90 | -30 | 53 | 14 |
| Anterior Insula |  |  | 12.20 | -30 | 23 | 2 |
| Middle Cingulate Cortex |  |  | 11.88 | 9 | 17 | 35 |
| Thalamus |  |  | 11.18 | 18 | -7 | -1 |
| Hippocampus |  |  | 10.65 | -27 | -25 | -7 |
| Parahippocampus |  |  | 9.84 | -21 | -28 | -10 |
| R. Anterior Insula | 13/45/47 | 1248 | 13.47 | 42 | 11 | -1 |
| Putamen |  |  | 12.17 | 24 | -1 | 11 |
| Thalamus |  |  | 11.18 | 18 | -7 | -1 |
| R. Anterior Prefrontal Cortex | 10 | 161 | 11.23 | 30 | 50 | 14 |
| R. Middle Frontal Gyrus | 6/8 | 176 | 9.66 | 36 | 2 | 59 |
| Precentral Gyrus |  |  | 9.61 | 48 | 8 | 47 |
| R. Middle Temporal Gyrus |  | 67 | 8.95 | 45 | -31 | -1 |
| L. Superior Temporal Gyrus |  | 28 | 8.88 | -57 | -46 | 20 |
| L. Middle Temporal Gyrus |  | 92 | 8.52 | -54 | -40 | -4 |
| L. Angular Gyrus | 7 | 115 | 8.16 | -27 | -55 | 32 |
| Superior Parietal Lobule |  |  | 7.10 | -30 | -67 | 47 |
| Precuneus |  |  | 6.59 | -12 | -55 | 29 |
| R. Hippocampus |  | 33 | 7.96 | 27 | -25 | -4 |
| L. Middle Temporal Gyrus |  | 32 | 7.84 | -54 | -10 | -16 |

Note that all regions are reported with a *p*_FWE_<0.001 threshold at the whole-brain level. L indicates left; R indicates right.
